# Coordinated CAII and CAIV establish a dual pH-regulatory axis essential for CatSper activation during sperm maturation and capacitation

**DOI:** 10.1101/2025.10.13.681600

**Authors:** Caroline Wiesehöfer, Cameron C. Gardner, Aura Stoskus, M. Agustina Battistone, Gunther Wennemuth, Jean-Ju Chung

## Abstract

Carbonic anhydrase II (CAII) and IV (CAIV) orchestrate pH regulation essential for male fertility through distinct cellular compartments, with CAII functioning intracellularly and CAIV GPI-anchored extracellularly in epididymal and vas deferens epithelia as well as on spermatozoa. These enzymes control pH environments required for sperm maturation and capacitation by regulating bicarbonate availability, which triggers soluble adenylyl cyclase/cAMP signaling, elevates intracellular pH (pH_i_), and enables motility activation. However, the specific contributions of CAII and CAIV to luminal pH and sperm pH_i_ regulation remain unclear. Here, we show that genetic ablation of either CAII or CAIV in mice impairs normal luminal acidification in the male reproductive tract and unexpectedly lowers the basal pH_i_ of sperm. This lower basal pH_i_ blunts subsequent intracellular alkalinization, thereby diminishing the activation of pH-sensitive CatSper Ca^2+^ channels essential for hyperactivated motility. Mechanistically, we demonstrate that this basal pH_i_ reduction occurs independently of the CatSper channel itself, but it is phenocopied in K^+^channel *Slo3*-deficient sperm, suggesting a homeostatic link between flagellar pH and membrane potential. Super-resolution imaging reveals that CAII is located within the flagellum near the CatSper channel in the principal piece, while CAIV is distributed along the entire flagellar membrane. Together, our findings demonstrate that CAII and CAIV employ spatially coordinated mechanisms to regulate pH. By maintaining an acidic luminal environment for sperm maturation and priming the subsequent alkalinization of sperm pH_i_, these enzymes modulate downstream ion channel activity to govern sperm motility and fertility.

**One-sentence summary:** Spatially coordinated carbonic anhydrases II and IV govern sperm motility by establishing the dual-compartment pH environments necessary to activate downstream flagellar ion channels.

## INTRODUCTION

Spermatozoa, after differentiation in the testis, must undergo further maturation processes in the male and female reproductive tracts to gain fertility competence ^1^. Luminal pH and bicarbonate (HCO_3_^-^) homeostasis throughout the male and female reproductive tracts are crucial to the entire reproductive process ^2,3^. The fluid in the epididymis, where sperm mature and are stored, is maintained as a slightly acidic environment with a very low HCO_3_^-^ concentration to sustain sperm viability and a quiescent state ^4–6^. In contrast, seminal plasma provides a more basic pH of 7.2–8.4 ^6,7^ and a high concentration of HCO_3_^-^, which not only neutralizes vaginal acidity but also begins to activate sperm motility ^3,8^. This early activation is further stimulated in the more alkaline, HCO_3_^-^-rich fluid of the uterus and oviduct ^9–11^.

Carbonic anhydrases (CAs) are crucial regulators of pH and HCO_3_^-^ production in the reproductive tracts and spermatozoa^12–14^, with CAII and CAIV being as among the most abundantly expressed proteins in human sperm ^14^. They reversibly catalyze the hydration of CO_2_ to HCO_3_^-^ and proton (CO_2_ + H_2_O ↔ HCO_3_^-^ + H^+^) ^15,16^, a reaction fundamental to pH homeostasis ^17^. CAs work in concert with other H^+^ and HCO_3_^-^ transport mechanisms in the epithelial cells of the reproductive tracts and in spermatozoa ^17–19^. More than a dozen CA isoforms have been identified in mammals, with CAII, CAIV, CAVII, and CAIX playing significant roles in CO_2_ hydration ^20^. CAII ^15^, CAIV ^21,22^, and CAXII ^23^ are robustly expressed in the epithelial cells of the epididymis and vas deferens, as well as in spermatozoa ^3,21,23,24^. Knockout mouse models of CAII and/or CAIV show reduced male fertility, highlighting the critical importance of these two isoforms in male reproduction ^22,24^. Spermatozoa from CAII and CAIV knockout mice exhibit an imbalance in HCO_3_^-^ homeostasis, resulting in decreased motility, including reduced swimming speed and beat frequency ^24^. Total CA activity in mouse spermatozoa remains consistent regardless of capacitation status, with CAII specifically accounting for half of the activity in capacitated spermatozoa ^24,25^.

The high extracellular HCO_3_^-^ allows sodium bicarbonate cotransporters (NBCs), specifically the solute carrier family SLC4, to bring HCO_3_^-^ into sperm cells ^26,27^. CAIV, as a GPI-anchored extracellular enzyme, rapidly converts extracellular CO_2_ into HCO_3_^-^ ^22,28,29^. This action increases the local HCO_3_^-^ concentration at the sperm surface, thereby facilitating its entry into the cell. Once inside, CAII helps to equilibrate the CO_2_/HCO_3_^-^ system ^24^, driven by concentration gradients ^30,31^. An increase in intracellular HCO_3_^-^ triggers sperm signaling during capacitation by activating sAC ^32^, thereby stimulating cAMP-dependent PKA activity ^33,34^. Additionally, HCO_3_^-^ raises sperm pH_i_ ^35,36^, activating the intracellular Ca^2+^ and pH-sensitive sperm-specific Slo3 K^+^ channel ^37–41^ and CatSper Ca^2+^ channel ^42–45^, both of which are essential for capacitation-induced sperm signaling ^46–50^. Despite this understanding, it remains unknown how CAII and CAIV coordinate their actions to maintain the acidic luminal environment in the epididymis and how they specifically contribute to local flagellar pH_i_ changes to regulate sperm ion channel activity.

Here, using *Car2* and *Car4* knockout mouse models, we demonstrate that CAII and CAIV are critical for acidifying the luminal pH of the cauda epididymis and vas deferens. Counterintuitively, despite the elevated luminal pH in these knockout mice, live sperm pH imaging reveals that both *Car2* and *Car4*-null spermatozoa exhibit a decreased basal pH_i_, impairing the alkalization needed for CatSper activation. Consistently, motility analyses of *Car2* and *Car4*-null spermatozoa show dysregulation of hyperactivation, supporting defective capacitation-associated Ca^2+^ signaling. Importantly, we show this reduced basal pH_i_ is not a consequence of CatSper dysfunction; rather, it is a primary feature shared by *Slo3*-null sperm. Using super-resolution microscopy, we visualize that CAII is specifically localized to the CatSper-containing principal piece, whereas CAIV is distributed throughout the entire flagellar membrane. Together, these findings reveal the coordinated yet compartment-specific mechanisms of CAII and CAIV in controlling sperm maturation and fertility. We establish the translational importance of this pH-regulatory axis by demonstrating that its pharmacological inhibition potently blocks hyperactivation in human sperm, identifying a fundamental mechanism conserved from mouse to man.

## RESULTS

### Genetic ablation of CAII or CAIV in mice impairs luminal acidification in the epididymis and vas deferens

To determine the role of CAII and CAIV in regulating luminal pH, we collected fluid from the cauda epididymis and vas deferens of anesthetized wild-type (wt), *Car2^-/-^*, and *Car4^-/-^* mice and immediately measured its pH using pH paper strips ^5,51^. In wt mice, the luminal fluid was acidic, with a pH of 6.4±0.1 in the cauda epididymis (**Fig. 1A**) and 6.6±0.1 in the vas deferens (**Fig. 1B**), consistent with previously reported values ^5,7,51^. Genetic ablation of either CAII or CAIV resulted in significant luminal alkalinization. In the cauda epididymis, the pH rose to 6.8±0.3 in *Car2^-/-^* mice and 6.6±0.3 in *Car4^-/-^* mice (**Fig. 1A**). Similar increases were observed in the vas deferens, where pH reached 6.8±0.2 in both knockout models (**Fig. 1B**). The pH shift was more pronounced in the absence of CAII (ΔpH=0.4) compared to CAIV loss (ΔpH=0.2) in the cauda epididymis, suggesting CAII plays a dominant role in luminal acidification. Despite these increases, the normal physiological gradient, lower in the cauda epididymis than the vas deferens, persisted across all genotypes. These results demonstrate that both intracellular CAII and GPI-anchored extracellular CAIV are essential for establishing and maintaining the optimal acidic luminal environment required for proper sperm maturation in the male reproductive tract.

**Figure 1.**
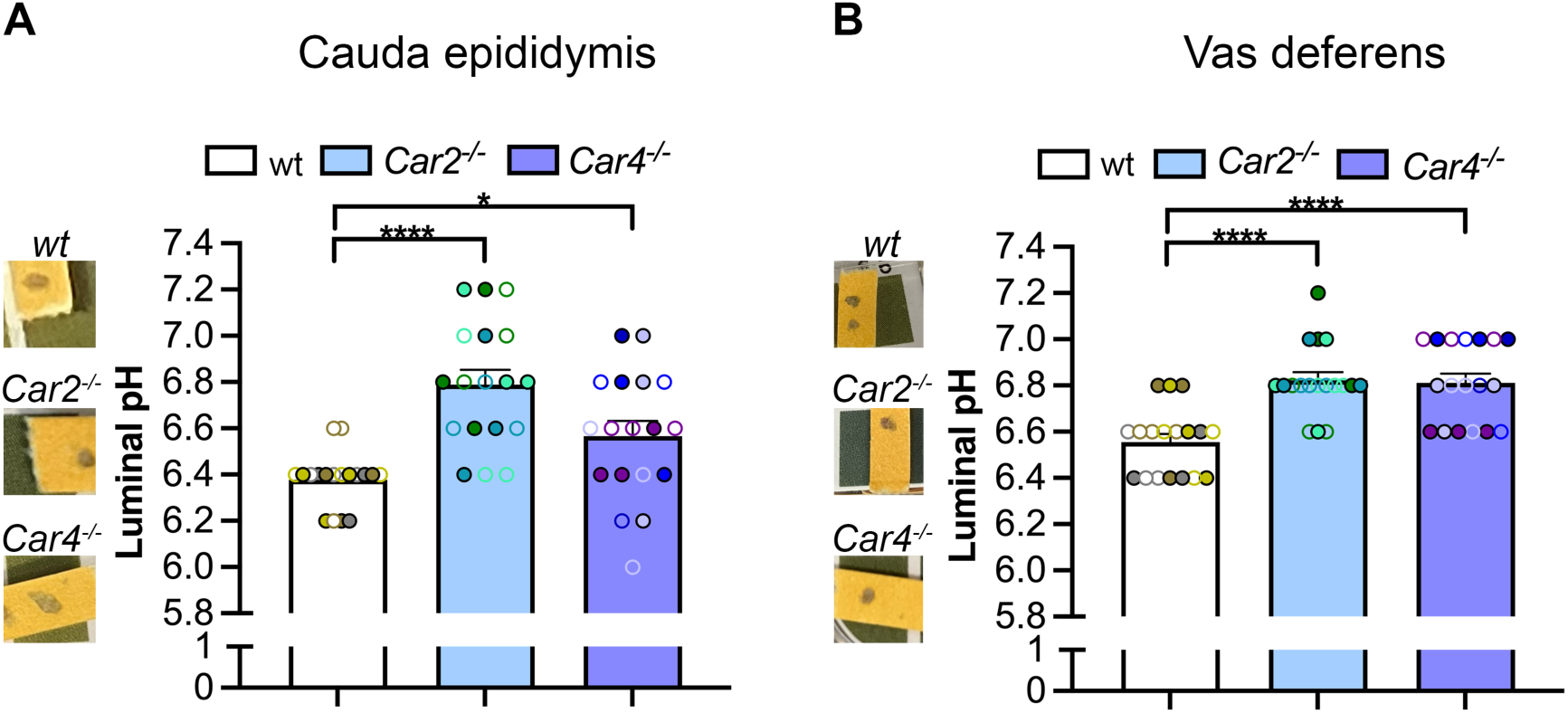
Impaired luminal acidification in the epididymis and vas deferens of *Car2^-/-^* or *Car4^-/-^*mice. (A and B) Luminal pH measurements from the cauda epididymis (A) and vas deferens (B) of wt (white), *Car2^-/-^* (blue), and *Car4^-/-^* (violet) mice. Each bar represents the mean ± SD from three independent animals (N=3) per genotype. A total of eighteen pH measurements were performed per genotype using both sides of the organs from each mouse with three technical replicates per organ. Statistical significance was determined by unpaired Students t-test. \**p* ≤ 0.05, ****p ≤ 0.0001. *See also* Supplemental Fig. 5.

### Carbonic anhydrase deficiency lowers basal sperm pH_i_ despite luminal alkalinization and impairs subsequent alkalinization

To assess the roles of CAII and CAIV in sperm pH_i_, we performed single-cell (**Fig. 2, Supplemental Fig. 1, A-G**) and population (**Supplemental Fig. 1, H-J**) pH analysis of motile spermatozoa isolated from the cauda epididymis and vas deferens using the fluorescent indicator, pHrodo-Red ^14,52,53^ (**Supplemental Fig. 1, A-C**). In non-capacitated wt sperm from the cauda epididymis, the basal pH_i_ was ∼6.4, closely matching the acidic luminal pH (**Fig. 1A, 2A and B**). Unexpectedly, despite the more alkaline luminal environment in knockout mice, epididymal sperm from *Car2*^-/-^ and *Car4*^-/-^mice exhibited dramatically lower basal pH_i_ values of ∼5.6 and ∼6.0, respectively (**Fig. 2A and B, Supplemental Fig. 1. D-E,** *see also* **Supplemental Fig. 5**).

**Figure 2.**
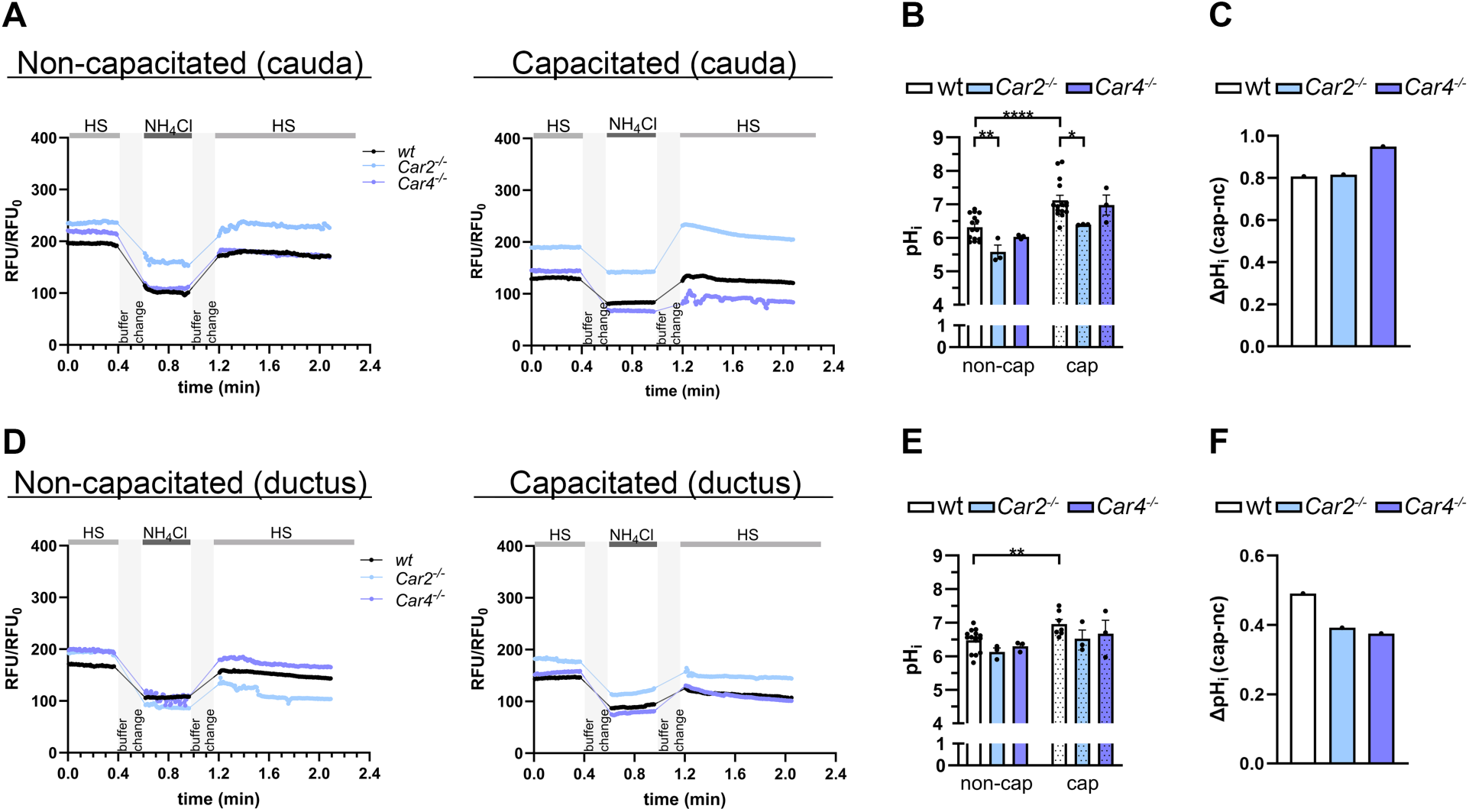
Carbonic anhydrases deficiency paradoxically lowers basal sperm pH_i_ and impairs pH_i_ recovery from an alkaline load. (A, D) Representative traces of pH_i_ change in sperm from cauda epididymis (A) and vas deferens (D). Sperm from wt (black), *Car2^-/-^* (blue) and *Car4^-/-^* (violet) mice were analyzed in non-capacitated (left panels) and capacitated (right panels) states before, during, and after exposure to 15 mM NH_4_Cl. Grey bars indicate buffer changes. (B, E) Quantification of basal (non-capacitated, open bars) and capacitated (dotted bars) pH_i_ in sperm from the cauda epididymis (B) and vas deferens (E), derived from traces as shown in (A) and (D). (C, F) Net pH_i_ increase (ΔpH_i_) during capacitation for sperm from the cauda epididymis (C) and vas deferens (F). ΔpH_i_ was calculated for each genotype by subtracting the mean non-capacitated pH_i_ from the mean capacitated pH_i_ shown in panels B and E. Data in B and E are shown as mean ± SEM. Sample sizes are indicated as N = number of mice and n= number of individual cells analyzed. Cauda sperm: Non-capacitated N_wt_=15, N_KOs_=3, n≥21, capacitated N_wt_=15, N_KOs_=3, n≥14. Vas deferens sperm: Non-capacitated N_wt_=15, N_KOs_=3, n≥18, capacitated N_wt_=7, N_KOs_=3, n≥14. Statistical significance was determined by unpaired Students t-test. *p ≤ 0.05, **p ≤ 0.01, ****p ≤ 0.0001. *See also* Supplemental Fig. 1 and 5.

Upon incubating under capacitation conditions, sperm from all genotypes alkalinized, indicating that the core pH_i_ regulatory machinery remains functional. This resulted in a comparable net pH_i_ increase during capacitation across all genotypes (**Fig. 2C**, cap ΔpH_i_ cauda: ∼0.8 in wt sperm, ∼0.8 in *Car2*^-/-^ sperm, and ∼0.9 in *Car4*^-/-^ sperm). However, the profoundly low starting pH_i_ of *Car2*^-/-^sperm meant its final pH_i_ reached only ∼6.4, a value similar to that of uncapacitated wt sperm and significantly lower than pH_i_ of ∼7.1 achieved by capacitated wt and *Car4*^-/-^ sperm (**Fig. 2B**). This dysregulation persisted in sperm from vas deferens (**Fig. 2D and E, Supplemental Fig. 1F-G**). Consistent with the higher luminal pH of this region, basal pH_i_ was slightly higher across all genotypes compared to cauda sperm (**Fig. 2B vs. 2E,** ΔpH_i_: ∼0.2 (wt), ∼0.6 (*Car2*^-/-^), and ∼0.3 (*Car4*^-/-^) sperm). Nonetheless, basal pH_i_ remained lower in sperm from *Car2*^-/-^ mice (∼6.1) compared to wt mice (∼6.5) (**Fig. 2E**). Capacitating conditions resulted in an intracellular alkalinization in all genotypes (**Fig. 2F**, cap ΔpH_i_: ∼0.5 (wt), ∼0.4 (*Car2*^-/-^), and ∼0.4 (*Car4*^-/-^) ductus sperm). These single-cell measurements were consistent with population-level analyses (**Supplemental Fig. S1, H and I**).

To directly assess the capacity of sperm to handle an alkaline load, we challenged them with a 15 mM NH_4_Cl addition (**Fig. 2, A and D**, **Supplemental Fig. S1, E and G**). Wt cauda sperm responded robustly, exhibiting a rapid pH_i_ increase of approximately 1.2 units. The response of *Car4^-/-^* sperm (ΔpH_i_ ∼1.4) was comparable to wt, indicating that extracellular CAIV is not essential for managing this type of internal alkaline challenge. In stark contrast, *Car2*^-/-^ cauda sperm displayed a significantly blunted response, with their pH_i_ increasing by only 1.0 unit. This impairment was even more pronounced after incubation under capacitating conditions (**Fig. 2, A and D, Supplemental Fig. S1, E and G**), highlighting the critical role of CAII when the cell’s buffering systems are under increased metabolic demand. Population-level analyses corroborated these single-cell measurements, including the attenuated alkaline response in *Car2*^-/-^ sperm (**Supplemental Fig. S1J**). These results demonstrate that while both CAs contribute to establishing basal pH_i_, intracellular CAII is uniquely critical for buffering rapid intracellular alkalinization for pH homeostasis.

### CA deficiency impairs CatSper activation and prevents sperm hyperactivation

Given the proposed unbalanced intracellular HCO_3_^-^ homeostasis in *Car2^-/-^*and *Car4^-/-^* sperm ^22,24^ and the significantly lower basal pH_i_ (**Fig. 2**), we next investigated the critical downstream consequence: the pH-dependent activation of the CatSper channel. We monitored intracellular [Ca^2+^] using Fluo4-AM and triggered CatSper opening by depolarizing alkaline solution (K8.6). Basal Ca^2+^ level in sperm from both knockout lines remains comparable to wt sperm (**Fig. 3, A and B**). However, *Car2^-/-^* sperm exhibited a significantly blunted Ca^2+^ response upon depolarization compared to wt sperm, with the defect being less pronounced in *Car4^-/-^* sperm (**Fig. 3, A and C**; ∼0.6-and 0.4-fold of wt, respectively). This suggests that the failure to establish a normal pH_i_ prevents CatSper from achieving full activation. Crucially, however, this impairment does not result in the profound signaling deficit observed in complete CatSper-null models ^54–57^. Unlike the aberrant capacitation-associated protein tyrosine phosphorylation (P-Tyr) potentiation seen in *Catsper1*-null sperm^54,55^ or following extracellular Ca^2+^ chelation ^58^, P-Tyr upregulation is preserved in both Car2^-/-^ and Car4^-/-^ sperm (**Fig. 3D**). Given that basal Ca^2+^ level remains unaffected (**Fig. 3, A and B**) and that the channels retain responsiveness to K8.6, this indicates that the Ca^2+^ influx in *Car* knockouts, while sub-maximal, remains within a range sufficient to maintain the signaling necessary for normal P-Tyr levels. Consistently, CATSPER1 protein levels remain unaltered in sperm from both *Car* knockouts (*see* **Supplemental Fig. 3F**).

**Fig 3.**
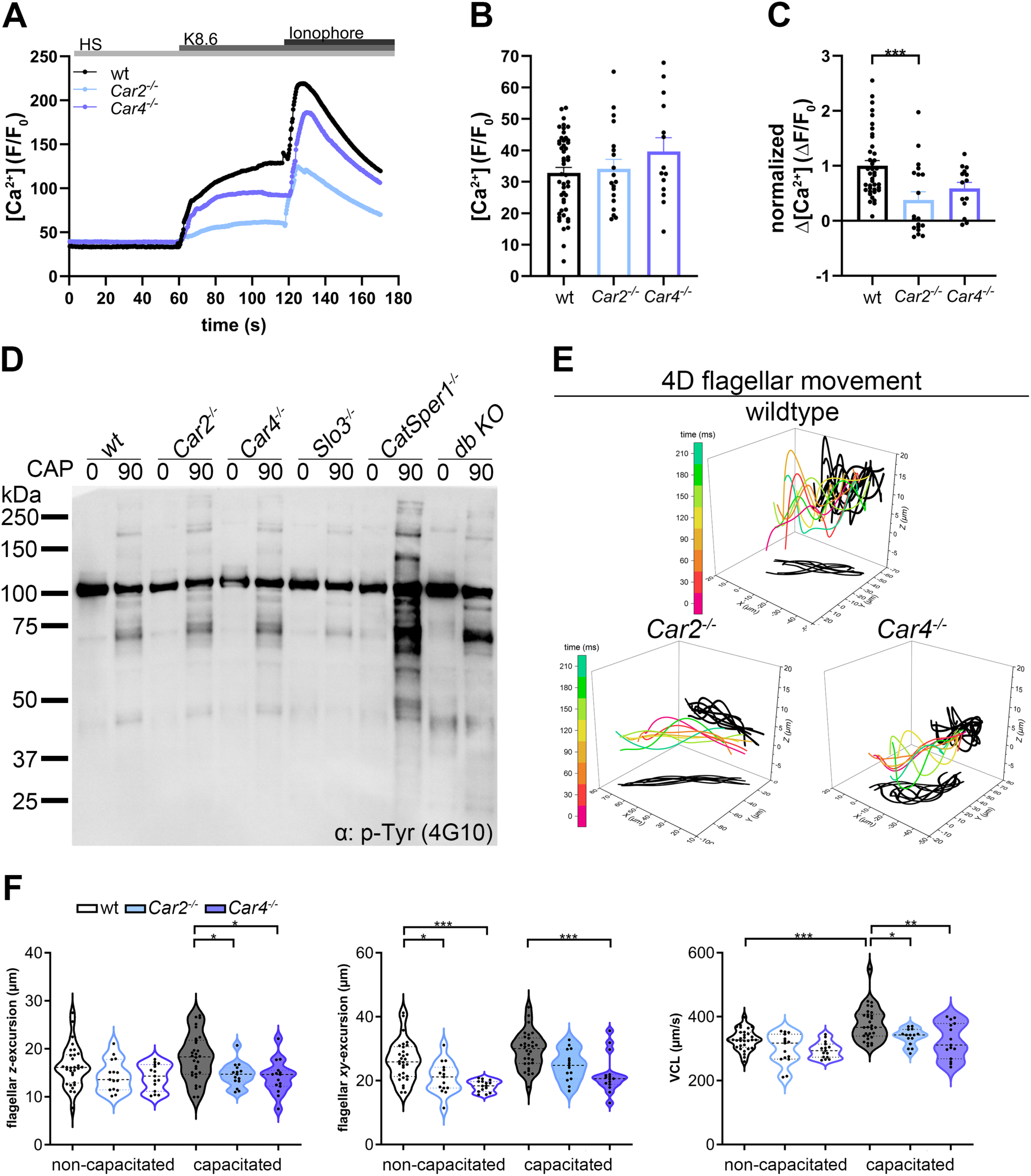
Genetic deletion of CAII or CAIV impairs CatSper-mediated Ca^2+^ influx and hyperactivated motility. (A-C) Intracellular Ca^2+^ responses in sperm from wt, *Car2^-/-^*, and *Car4^-/-^* mice. Sperm were loaded with the Ca^2+^ indicator Fluo4-AM and stimulated with high-potassium medium (K8.6) to trigger CatSper-dependent Ca^2+^ influx. (A) Representative mean fluorescence traces (F/F_0_) over time in sperm from wt (black), *Car2^-/-^* (blue), and *Car4^-/-^* (violet) mice. (B) Basal Ca^2+^ level determination of sperm analyzed in A over 58 s. (C) Quantification of the peak Ca^2+^ change (ΔF/F_0_), normalized to the wt response over 58 s. The Ca^2+^ ionophore ionomycin (5 µM) was used as a positive control to determine maximum fluorescence. Data are from N≥4 mice, with n≥15 cells. (D) Immunoblot analysis of protein tyrosine phosphorylation (pTyr). Protein lysates were collected from sperm populations before (0 min) and after (90 min) incubation under capacitating conditions. (E) Representative 4D flagellar trace of a single sperm analyzed by DHM. The color-coded trace shows the flagellar excursion over time (0-210 ms), with black shadows representing its projections onto the XY and XZ planes, illustrating the complex 3D beat pattern. (F) Quantification of key 4D motility parameters in non-capacitated (open) and capacitated (filled) sperm in violin plots. Each point represents a single sperm (*n*> 14 per condition). Data are presented showing the median (thick dash lines) and interquartile range (thin dash lines). Asterisks indicate a significant difference from wt under the same condition: \**p* ≤ 0.05, \*\**p* ≤ 0.01, \*\*\**p* ≤ 0.001. *See also* Supplemental Fig. 2.

While this sub-maximal Ca^2+^ response preserves basal signaling, it is clearly insufficient to sustain the normal flagellar motility. To determine how these molecular bottlenecks manifest in physical behavior, we assessed flagellar dynamics using both standard CASA and 4D motility analysis using Digital Holographic microscopy (DHM) ^59–61^. Consistent with previous results ^22,24^, total sperm count was reduced in the KO males (**Supplemental Fig. 2A**). Initial 2D CASA analysis revealed that total motility was reduced 4.5-fold in *Car2^-/-^* sperm and 1.6-fold in *Car4^-/-^* sperm (**Supplemental Fig. 2B**). 4D motility analysis further revealed that, under non-capacitating conditions, *KO* sperm showed altered planar (*XY*) flagellar excursions but normal vertical (*Z*) excursions (**Fig. 3, E and F**). More crucially, the ability to transition to a hyperactivated state during capacitation was severely impaired. Both *Car2*^-/-^ and *Car4^-/-^* sperm failed to develop the high-amplitude, asymmetric flagellar beats characteristic of hyperactivation. This was quantified as a significant reduction in flagellar excursion in both *XY* and *Z* planes, curvilinear velocity (VCL) and lateral head excursion (**Fig. 3E and F**; **Supplemental Fig. 2C**). Taken together, these data indicate that the lower basal pH_i_ in CA-deficient sperm compromises pH-dependent CatSper activation, leading to defective hyperactivation. This functional impairment is particularly severe in *Car2*^-/-^ sperm, aligning with its more pronounced pH_i_ and Ca^2+^ response defects.

### Loss of SLO3 impairs sperm pH homeostasis in mice

Having established that CA deficiency impairs pH_i_ and subsequent Ca^2+^ influx, we next investigated the molecular interplay between CAs, the CatSper complex, and the K^+^ channel SLO3−another major regulator of sperm capacitation. We first measured pH_i_ in sperm lacking these channels (**Fig. 4, A-D**). *Catsper1^-/-^* sperm, which lack the entire channel complex, exhibited pH_i_ values statistically indistinguishable from wt sperm at both basal and capacitating conditions (**Fig. 4, A-D, Supplemental Fig. 3, A and B**). This indicates that CatSper-mediated Ca^2+^ entry and its downstream signaling are not primary determinants of sperm pH_i_ regulation.

**Figure 4.**
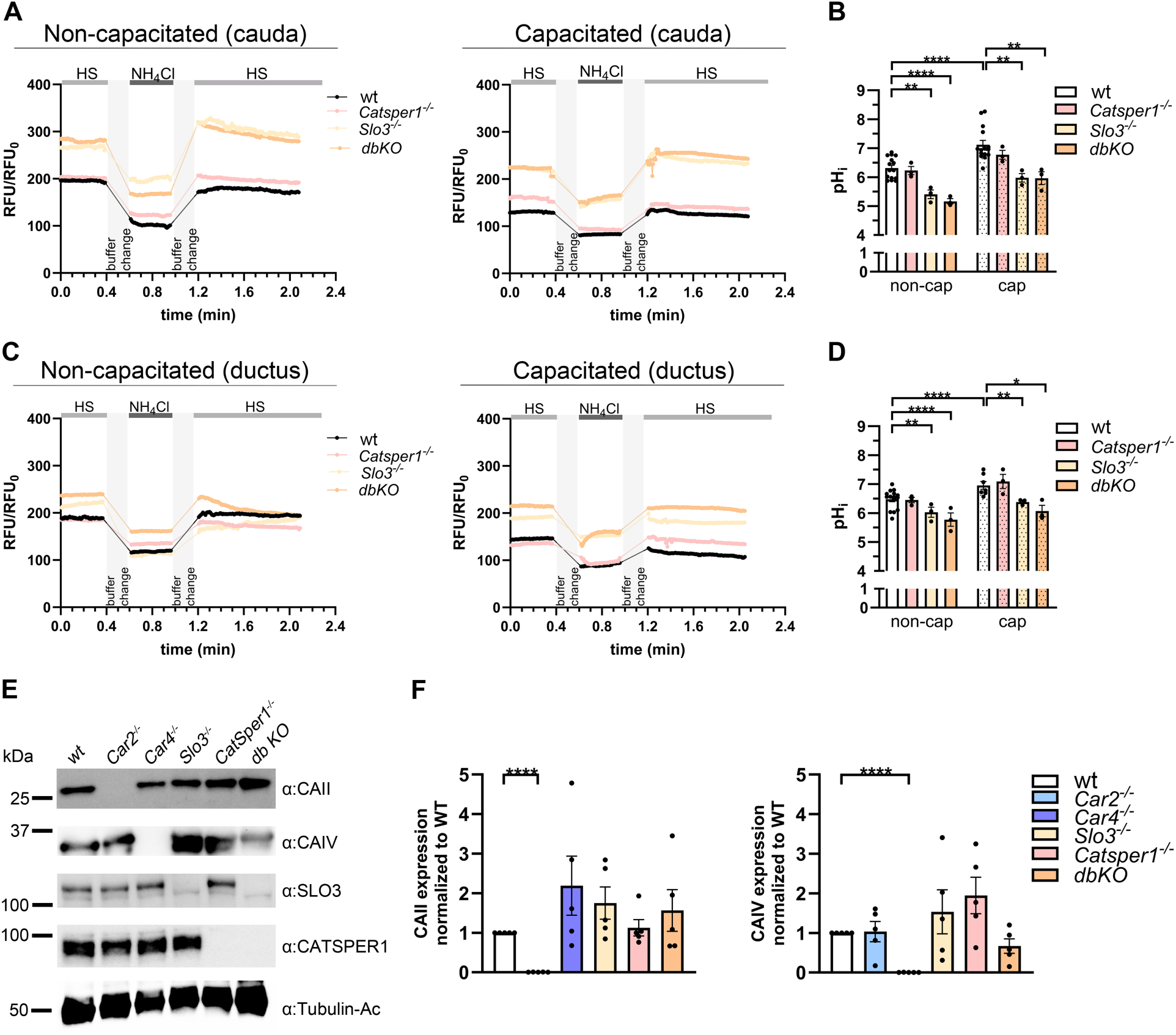
**Loss of the SLO3 channel disrupts pH_i_ homeostasis**. (A-D) Analysis of intracellular pH (pH_i_) in cauda epididymal sperm. (A) Representative live-cell pH_i_ traces from sperm loaded with pHrodo Red-AM. Sperm from wt (black), *Catsper1^-/-^* (pink), *Slo3^-/-^* (beige), and *Slo3^-/-^, Catsper1^-/-^*double knock-out (*dbKO*, orange) mice were analyzed under non-capacitating conditions (left) and capacitating (right) conditions. Sperm were challenged to an alkaline load via a 15 mM NH_4_Cl stimulation (gray bar) to assess their pH_i_ handling capacity. (B) Quantification of basal pH_i_ of non-capacitated (open bars) and capacitated (dotted bars) sperm, calculated from fluorescence intensity and calibration curve from Supplemental Fig. 1C. (C-D) Analysis of pH_i_ in vas deferens sperm. (C) Representative pH_i_ traces of vas deferens sperm, performed as described in (A). (D) Quantification of basal pH_i_ of non-capacitated and capacitated vas deferens sperm. (E) Representative immunoblots showing protein levels of CAII and CAIV in sperm lysates from wt, *Car2^-/-^, Car4^-/-^, Slo3^-/-^*, *Catsper1^-/-^* and *dbKO* mice. (F) Densitometric quantification of CAII (*left graph*) and CAIV (*right graph*). Data in (B, D, F) are presented as mean ± SEM. Sample size for pH_i_ experiments: N=3-15 mice per genotype, n≥14-18 cells per experiment. For protein quantification (F), each point represents a biological replicate from an individual male. Asterisk indicates a significant difference from wt under the same condition: \**p* ≤ 0.05, \*\**p* ≤ 0.01, \*\*\*\**p* ≤ 0.0001. *See also* Supplemental Fig. 3.

In stark contrast, loss of SLO3 profoundly acidified sperm pH_i_. Both *Slo3^-/-^* and *Catsper1^-/-^;Slo3^-/-^*double-knockout sperm displayed significantly lower basal pH_i_ (∼5.4 and ∼5.2, respectively) compared to wt (∼6.4) (**Fig. 4, A and B**). This severe acidification persisted under capacitating conditions (∼6.0; and ∼6.2, respectively) and is consistent with recent reports linking membrane potential with pH_i_ regulatory function ^52,62^. This pH_i_ defect was observed in sperm from both the cauda epididymis (**Fig. 4, A and B)** and the vas deferens (**Fig. 4, C and D**) and was confirmed by population-level analysis and NH_4_Cl stimulation (**Supplemental Fig. 3, C-E**). The striking acidification of *Slo3*^-/-^ sperm, which phenocopies the defect observed in *Car2*^-/-^ and *Car4*^-/-^ mice, prompted us to examine whether this phenotype resulted from altered CA protein levels. However, CAII and CAIV protein levels are unchanged in sperm regardless of the absence or presence of CatSper or SLO3 (**Fig. 4, E and F; Supplemental Fig. 3F**). These findings demonstrate that the severe pH_i_ reduction in *Slo3*^-/-^ sperm is independent of CAII or CAIV protein abundance, suggesting that SLO3 and CA isoforms maintain pH_i_ through distinct, yet functionally convergent, pathways.

### Super-resolution microscopy reveals distinct flagellar domains for CAII and CAIV

To understand how CAII and CAIV might exert their distinct effects, we examined their subcellular localization in wt sperm. Confocal microscopy revealed different distribution patterns: CAII was highly enriched in the principal piece—the segment housing the CatSper channel ^44,54^—and the acrosomal region (**Fig. 5A**, *first row*) whereas CAIV was distributed along the entire length of the flagellum (**Fig. 5A**, *second row*).

**Figure 5.**
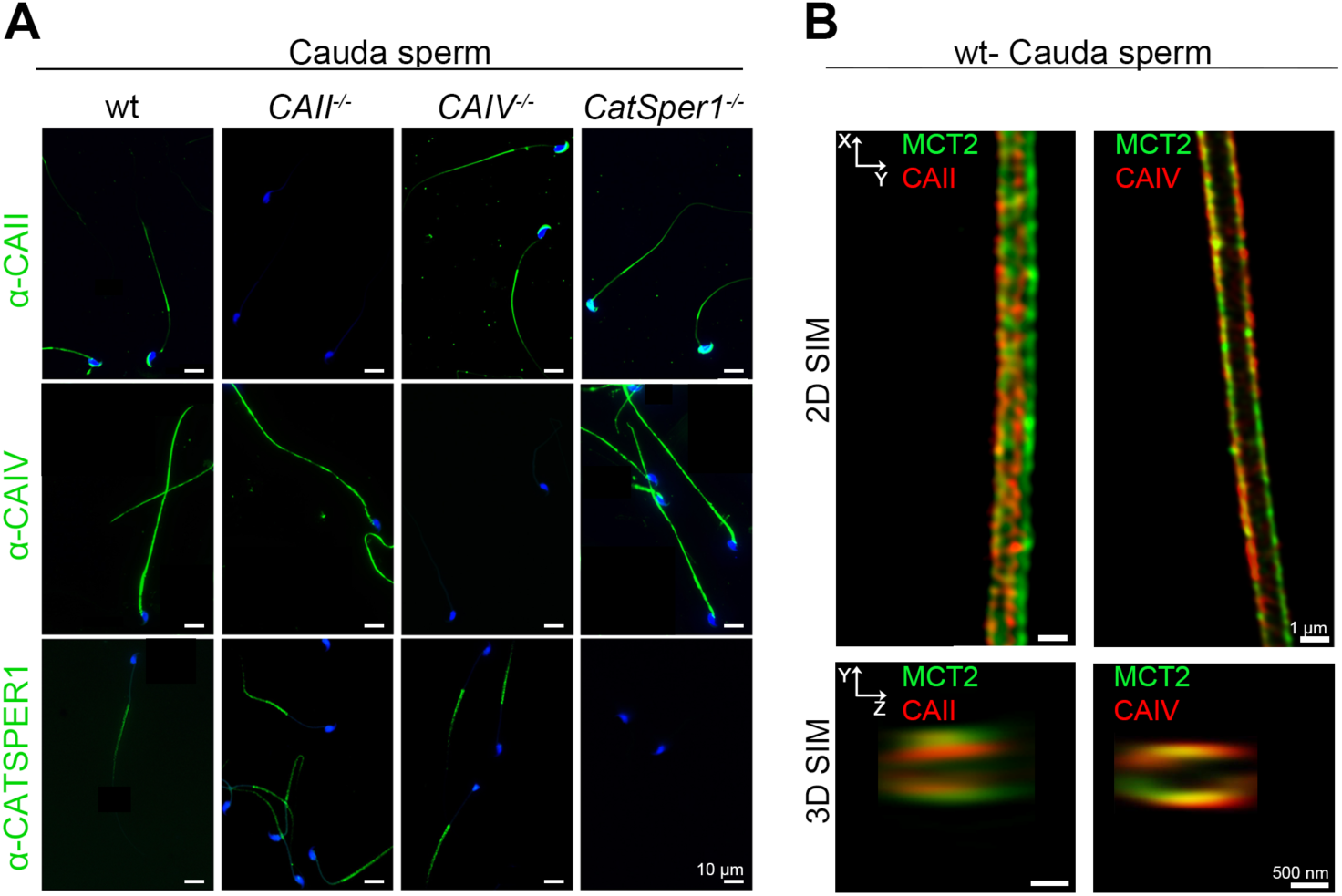
CAII and CAIV exhibit distinct subcellular localization within the sperm flagellum. (A) Confocal immunofluorescence microscopy of CAII, CAIV, and CATSPER1. Images show the localization of CAII (*top row*), CAIV (*middle row*) and CATSPER1 (*bottom row*) in sperm from wt and knockout animals. The specific knockout sperm (*Car2^-/-^*, *Car4^-/-^*, *Catsper1^-/-^*) serve as negative controls, confirming specificity. (B) Super-resolution 3D-SIM imaging reveals the proximity of CAs to the flagellar membrane. High-resolution images showing the intraflagellar localization of CAII (*left panels*) and CAIV (*right panels*) relative to the monocarboxylate transporter 2 (MCT2), a known flagellar membrane marker. Top row: *XY*-projections show the view along the flagellum. Bottom row: *YZ*-projections show cross-sectional views. CAII and CAIV (red), MCT2 (green). *See also* Supplemental Fig. 4.

To achieve sub-diffraction resolution and define their positions relative to the flagellar membrane, we used structured illumination microscopy (SIM). This high-resolution imaging confirmed that the GPI-anchored protein CAIV ^63,64^ extensively co-localizes with at the flagellar membrane with monocarboxylate transporter 2 (MCT2), a well-characterized flagellar membrane transporter ^65–67^ (**Fig. 5B**; **Supplemental Fig. 4**). Conversely, while CAII is cytosolic, SIM confirmed its enrichment just beneath the flagellar membrane specifically within the principal piece, in close proximity to, yet distinct from, MCT2. Together, these data demonstrate that CAII and CAIV occupy spatially distinct domains: CAIV functions as a membrane-associated extracellular enzyme along the entire flagellum, while CAII is compartmentalized intracellularly within the principal piece. This architectural separation provides a clear compartmental basis for their differential contributions to local pH_i_ regulation and ion channel activity.

### Pharmacological inhibition of CAs impairs hyperactivation in both mouse and human sperm

To confirm our genetic findings and test the conservation of CA function, we treated both mouse and human sperm with the pan-carbonic anhydrase inhibitor ethoxzolamide (EZA) ^22,24^ and performed single-cell 4D motility analysis ^59,60^. In mouse sperm, EZA treatment specifically blocked the development of hyperactivated motility. While control sperm displayed the expected increase in VCL, ALH, and flagellar *XY*-excursion following incubation in capacitating medium, sperm treated with EZA failed to achieve this hyperactivated state (**Fig. 6A-C**).

**Fig 6.**
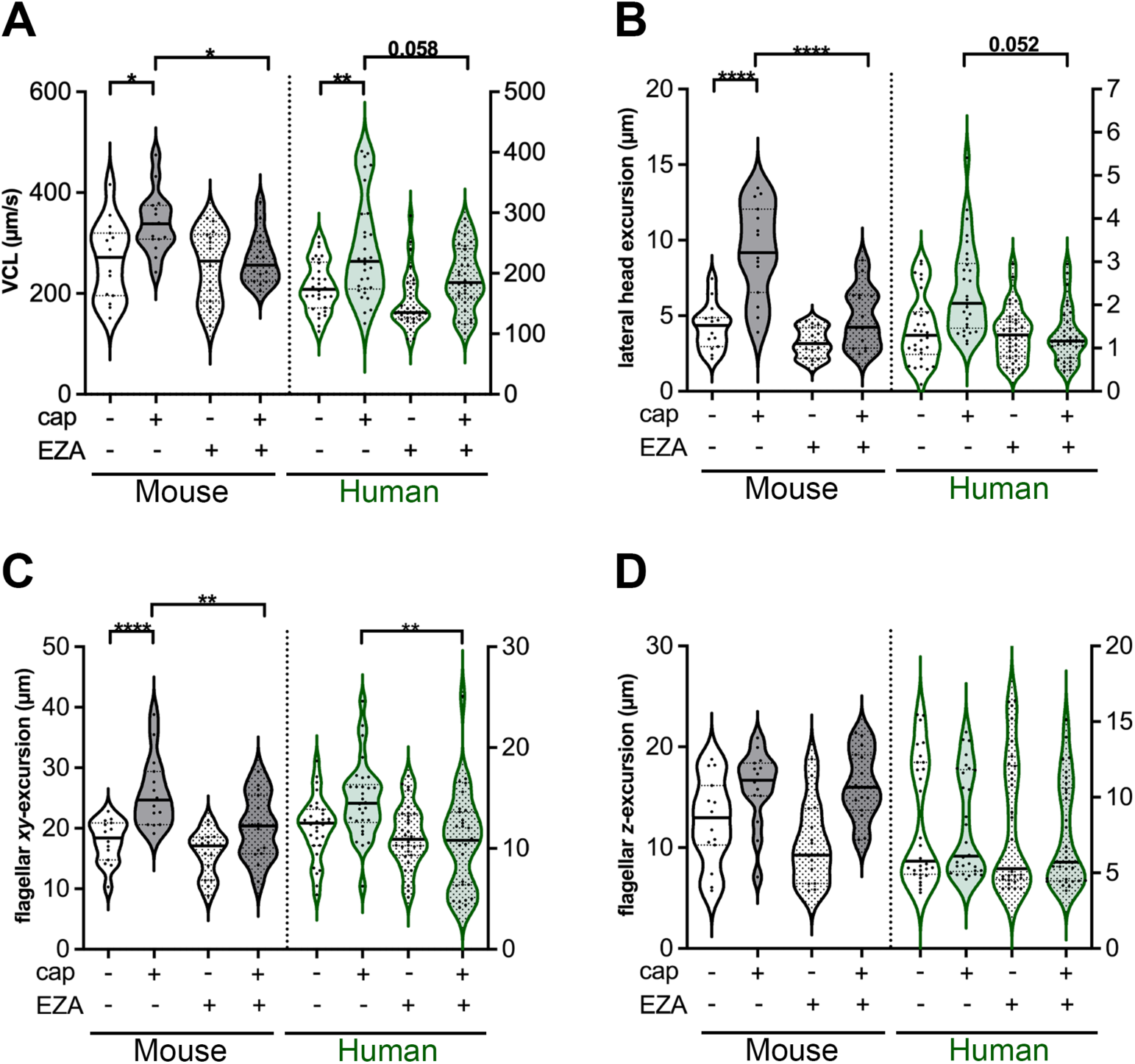
Pharmacological inhibition of CAs impairs hyperactivated motility in both mouse and human sperm. (A-D) 4D sperm motility analyzed using DHM. Sperm from mice and humans were incubated under non-capacitating or capacitating conditions in the presence of either vehicle (control) or the pan-carbonic anhydrase inhibitor ethoxyzolamide (EZA, 5 µM). The plots compare key motility parameters between control and EZA-treated groups. Data from mouse sperm are outlined in black, and human sperm in green. Non-capacitated conditions are represented by open plots, and capacitated conditions by filled plots. Quantifications of 3D curvilinear velocity (VCL) (A), Lateral head displacement (B), Flagellar planar excursion (*XY*-excursion) (C), and Flagellar vertical excursion (*Z*-excursion) (D). Each data point represents a single analyzed sperm (N_mouse_= 3, n_mouse_=15 per group; N_human_= 6; n_human_= 30 per group). Plots show median (thick dash line) and interquartile range (thin dash lines). Asterisks denote a significant difference between the control and EZA-treated group within the same condition: *p ≤ 0.05, **p ≤ 0.01, ***p ≤ 0.001, ****p ≤ 0.0001.

EZA also prevented hyperactivation in human sperm, indicating that this functional requirement is conserved. Under capacitating conditions, EZA impaired the increase in VCL (*p*= 0.058), head excursion (α= 0.052), and the flagellar *XY*-excursions significantly (**Fig. 6, A-C**). Although some species-specific nuances were observed−for instance, EZA reduced the basal VCL of non-capacitated human sperm, whereas flagellar *Z*-excursion remained unaffected in either species (**Fig. 6, A and D**). Computer-assisted sperm analysis (CASA) to measure sperm population also showed a reduction in VCL in capacitated human sperm due to EZA (s*ee* **Supplementary Fig. 2D**). These pharmacological data demonstrate that CA activity is required for achieving hyperactivated motility across mammalian species.

## DISCUSSION

### CAs are context-dependent pH regulators in male reproduction

In this study, we reveal that CAII and CAIV are critical for establishing the correct pH in two distinct compartments: the luminal environment of the male reproductive tract and, intrinsically, the sperm cell itself. A key insight from our work is that CAs operate via different mechanisms in these two locations. In the epididymal and vas deferens epithelia, where bicarbonate concentration is tightly regulated, CAII and CAIV appear to coordinate to maintain an acidic luminal environment (**Supplemental Fig. 5A**). In stark contrast, within the sperm, they act in concert to facilitate the generation of intracellular bicarbonate (HCO_3_^-^), which is essential for pH_i_ homeostasis and downstream signaling (**Supplemental Fig. 5B**). We demonstrate that disrupting this intrinsic pH regulation is the upstream cause of impaired CatSper channel activation.

Importantly, our data establish that this is a unidirectional pathway: genetic ablation of CatSper does not alter sperm pH_i_. This finding clarifies previous observations, such as those reporting a decreased basal pH_i_ in *CatSper1*-null sperm ^68^. We attribute this discrepancy to differences in experimental sensitivity; the high signal-to-noise ratio and fluorogenic properties of the pHRodo probe ^2,52,53^, when corroborated across both single-cell and population-level measurement, provide a robust foundation for our conclusion that sperm pH_i_ is regulated independently of CatSper-mediated Ca^2+^ entry. By decoupling these events, our work repositions the CA-mediated pH axis as a fundamental, upstream regulator of the CatSper complex, rather than a downstream consequence of it.

### Coordinated roles of CAII and CAIV in luminal acidification and sperm pH homeostasis

A key requirement for sperm maturation and storage is the maintenance of an acidic, low-bicarbonate fluid in the epididymis and vas deferens ^3,69^. Our results confirm that both CAII and CAIV are indispensable for this process, as genetic ablation of either enzyme leads to a significant rise in luminal pH (**Fig. 1**). This points to a sophisticated, coordinated mechanism for luminal pH regulation (**Supplemental Fig. 5A**). The profound pH increase in *Car2*^-/-^ mice supports the role of CAII as the primary intracellular “engine” for generating the protons subsequently secreted into the lumen by vacuolar H^+^-ATPases (V-ATPases) on the apical membrane of epididymal clear cells ^70–72^. A raised luminal pH in the absence of CAII is therefore a direct consequence of the loss of this proton fuel source. The significant, albeit less severe, pH rise in *Car4*^-/-^ mice highlights the crucial role of extracellular CAIV, anchored in the apical region of the principal cells in the corpus and cauda epididymides ^24,73^. We propose that CAIV facilitates scavenging luminal bicarbonate (H^+^ + HCO_3_^-^ → CO_2_ + H_2_O) ^17,73,74^. By reducing the concentration of this primary buffer, CAIV allows for more efficient H^+^ pumping by V-ATPases, while simultaneously generating CO_2_ that can diffuse back to the cell for use by CAII ^17^. In essence, CAII provides the protons, while CAIV ensures that these protons effectively lower the luminal pH by depleting the bicarbonate buffer.

While the epididymal lumen of CA-deficient mice was more alkaline, the sperm exhibited a significantly lower basal pH_i_ (**Fig. 2**). This critical finding suggests a cell-autonomous defect and a fundamentally different operational model for CAs within sperm (**Supplemental Fig. 5B**). We propose that in sperm, both CAs catalyze the same forward reaction (CO_2_ + H_2_O → H^+^ + HCO_3_^-^). When sperm encounter the alkaline fluid of the female reproductive tract, CAIV on the sperm surface converts extracellular CO_2_ into HCO_3_^-^, increasing the local concentration to facilitate its entry via transporters like solute carrier-family bicarbonate transporters ^11^. Internally, HCO_3_^-^, generated by both CAs, acts as the primary buffer against the protons produced during rapid glycolytic flux, the primary metabolic engine of mature sperm ^75,76^. Without efficient HCO_3_^-^production, sperm cannot effectively mitigate this acid load, resulting in a constitutively lower pH_i_.

Crucially, acute pharmacological inhibition of CAs phenocopies these motility defects in both mouse and human sperm (**Fig. 6**). demonstrating a direct CA role in mature sperm and a deeply conserved principle. Despite minor species-specific variations, such as the observed differences in basal VCL, the consistency of our findings across animal models underscores the evolutionary conservation of this regulatory axis. Together, these results demonstrate that CA activity is required for achieving hyperactivated motility across mammalian species, highlighting a deeply conserved principle in sperm physiology.

### Imbalanced HCO_3_^-^ and low basal pH_i_ are the causes of motility and CatSper defects in CA-deficient sperm

Disturbances in luminal pH are closely linked to sperm motility dysfunction ^51^, suggesting that luminal pH homeostasis is fundamental to sperm maturation. This motility failure, which our DHM analysis captures with unprecedented 3D detail, can be explained by the collapse in the core signaling cascades that govern capacitation. This cascade is initiated by HCO_3_^-^ ^77–79^, which acts as a crucial second messenger with two distinct but interconnected roles. First, HCO_3_^-^ directly stimulates soluble adenylyl cyclase (sAC) ^77,80^, leading to cAMP production and PKA activation--the key drivers of capacitation-associated tyrosine phosphorylation ^78,81^. Second, and central to our findings, the hyperactivation failure can also be directly traced to impaired CatSper channel activity (**Fig. 3**). CatSper is pH-sensitive and requires an alkaline pH_i_ for full potentiation ^42,43,82^. Therefore, one possible explanation for impaired CatSper channel activity is that the constitutively acidic intracellular environment of *Car2^-/-^* and *Car4^-/-^* sperm clamps CatSper in a low-activity state, preventing the Ca^2+^ influx needed for hyperactivation (**Fig. 3**, **Supplemental Fig. 5B**).

Such an attenuated activation of CatSper by pH_i_ acidification is consistent with the reduced Ca^2+^ sensitivity of the channel under acidic conditions ^43^. Furthermore, our data aligns with models suggesting that the pH_i_ regulatory network is interconnected: reduced SLO3 activation leads to insufficient hyperpolarization, which in turn diminishes the driving force for proton efflux via voltage-dependent mechanisms such as sNHE ^83,84^, evokes flagellar pH_i_ acidification ^52^, which also prevent CatSper from full activation. While our data highlights pH_i_ as a primary gating mechanism, the signaling network governing hyperactivation is complex. Interestingly, other studies have described a HCO_3_^-^ dependent modulation of Ca^2+^ entry via sAC-generated cAMP ^33,85,86^, further supporting the broader requirement of CAII and CAIV in driving the upstream regulation of CatSper regulation.

### CAII and CAIV are spatially organized and functionally coupled to ion transporters for pH_i_ control

In mouse sperm, the transition to hyperactivation is critically dependent on a pH_i_ increase from a basal level of ∼6.6 ^62,87^ to ∼7.0 ^87,88^ (**Fig. 2**). This pH_i_ rise, driven by HCO_3_^-^ uptake ^3,26,88^, acts as a permissive signal that directly gates key ion channels, including SLO3 and CatSper ^37,39,41,43^. The activation of SLO3 is a prerequisite for the membrane hyperpolarization necessary for capacitation ^39,40^, and the subsequent opening of the CatSper channel is the definitive trigger for the Ca^2+^ influx that drives hyperactivated motility ^44,46,84^. We propose that the primary defect in *Car2*^-/-^ and *Car4*^-/-^ sperm is a breakdown in this fundamental regulatory axis. By failing to efficiently produce intracellular HCO_3_^-^, the sperm cannot adequately buffer the constant acid load generated by its own metabolism—namely, protons from glycolysis in the principal piece and oxidative phosphorylation in the midpiece ^89,90^. This metabolic mismatch results in the constitutively low pH_i_ we observed, which prevents the sperm from reaching the pH threshold required to activate SLO3 and CatSper, ultimately explaining their severe motility deficits.

To explain their non-redundant, cell-autonomous roles, we propose that CAII and CAIV operate within a sophisticated “transport metabolon”, functionally coupling with distinct acid-base transporters ^19,91–95^. While direct biochemical evidence of a CA-metabolon in sperm is pending, the conceptual precedent is strong. Recent high-resolution cryo-electron microscopy structures—specifically the native CatSper complex architecture ^61^ and the sperm-specific radial spoke 3 embedded with kinases and metabolic enzymes ^96^—reveal that flagellar signaling relies on highly organized, nanoscale protein complexes. In other tissues, the direct physical association of CAs with transporters maximizes ion flux by rapidly supplying or removing substrates at the transporter pore, bypassing the limits of bulk diffusion ^19,93,97,98^. For example, the interaction between CAIV and Na^+^/HCO_3_^-^ co-transporter is known to drive rapid HCO_3_^-^ influx and alkalization in nasal epithelial cells ^99^.

In conclusion, the global loss of CAII or CAIV produces a complex, dual-compartment pH-dysregulation that affects both the luminal environment and the sperm-intrinsic functions critical for fertility. In the epididymis epithelia, these enzymes coordinate their efforts to acidify the lumen. In the sperm, they act in concert to generate the bicarbonate that governs pH_i_ homeostasis and the capacitation cascade, standing as the critical upstream gatekeepers for CatSper activation and the acquisition of fertilizing ability.

## MATERIALS and METHODS

### Antibodies

Mouse monoclonal anti-mouse CATSPER1 antibody was described previously ^57^. Rabbit polyclonal anti-mouse MCT2 antibody was generated in this study by immunizing rabbits with KLH-conjugated peptide (MEALNRSKQDEVTV-C). Antisera from the immunized rabbits were affinity-purified using the peptide immobilized SulfoLink coupling resin (Thermo Fisher). To validate anti-MCT2 antibody specificity, the anti-MCT2 antibody was preabsorbed with the antigenic peptide at a 1:10 weight-to-weight ratio for 1 hour at room temperature. All other antibodies used in this study are commercially available and are listed in Supplemental Table 1.

### Reagents

All reagents used in this study are listed in Supplemental Table 2.

### Animals

C57BL/6J wild-type, *Car2*- and *Car4*-null mice were obtained from Jackson Laboratories (strain numbers 000664, 001623, and 008217, respectively). *Catsper1^-/-^* and *Slo3^-/-^* mice were generated in previous studies ^41,44^. *Catsper1^-/-^*;*Slo3^-/-^* double knockout is generated by crossing the two lines. All mice were maintained on C57BL/6J background. Mice were managed and treated in accordance with the guidelines approved by the Yale Institute Animal Care and Use Committees (IACUC, #20079), Massachusetts General Hospital/Harvard Medical School (IACUC, #2003N000216) and University of Duisburg-Essen (LANUV, #AZ81-02.04.2020.A493).

### Murine sperm preparation

Mouse sperm were isolated from the cauda epididymis or vas deferens of 3-6 month old adult male mice by the swim-out method in EmbryoMax^®^ M2 medium (EMD Millipore) or modified HEPES buffered saline (HS) solution containing (in mM: 135 NaCl, 5 KCl, 2 CaCl_2_, 1 MgCl_2_, 20 HEPES, 5 glucose, 10 lactic acid and 10 pyruvic acid, pH 7.4, 353 mOsm). After two centrifugation steps (3 min, 500 g), the spermatozoa were resuspended in regular HS buffer. *In vitro* capacitation was induced by incubating 2×10^6^ sperm/ml in capacitation medium, either in HS supplemented with 15 mM HCO_3_^-^ + 5 mg/ml BSA, pH 7.4, 374 mOsm or human tubular fluid (HTF) medium (EMD Millipore) at 37°C, 5% CO_2_ for 90 min. For pH_i_ analysis, capacitated spermatozoa were washed once and measured in the regular HS buffer without bicarbonate supplement.

### Human sperm preparation

A panel of three donors provided ejaculated human sperm after a sexual abstinence of 72h. By the use of a swim-up procedure as previously described ^60^, human sperm were separated from the ejaculates. In short, 0.5 ml of the sperm suspension was layered under 2 ml of modified HEPES buffered saline (HS) solution containing (in mM: 135 NaCl, 5 KCl, 2 CaCl_2_, 1 MgCl_2_, 20 HEPES, 5 glucose, 10 lactic acid and 10 pyruvic acid, pH 7.4, 353 mOsm). After 60 min incubation at 37 C and 5% CO_2_ supernatant containing motile sperm was used for further experiments. *In vitro* capacitation was induced by incubating 2×10^6^ sperm/ml in capacitation medium (HS supplemented with 15 mM HCO_3_^-^ + 5 mg/ml BSA, pH 7.4, 374 mOsm) at 37°C, 5% CO_2_ for 4 hours. All experiments with human spermatozoa were approved by the Ethics Committee of the University of Duisburg-Essen (14-5748-BO and Addendum 11/2019) and the samples were de-identified with all donors signed informed consent forms prior to sample donation.

### Sperm protein isolation and Western Blot analysis

Whole sperm protein content was extracted as previously described ^54,100^. Briefly, mouse epididymal spermatozoa were washed once in PBS and lysed in 2X LDS sample buffer (GenScript). The solubilized sperm lysate collected after centrifugation at 15,000 g, 4°C for 10 min was adjusted to 50 mM DTT and denatured at 75°C for 10 min before loading onto the gel. For Western blot analysis, mouse anti-CATSPER1 (1:1,000), sheep anti-CAII (1:2,000), goat anti-CAIV (1:1,000), mouse anti-SLO3 (1:1,000), mouse anti-pTyr (4G10) (1:2,000), and mouse anti-acetylated tubulin (1:20,000) were used for primary antibodies. Corresponding secondary antibodies (Jackson ImmunoResearch) for the species were used in 1:10,000 dilution. Protein bands were visualized using Clarity™ Western ECL Substrate or Pierce^TM^ ECL Western blotting substrate and ChemiDoc™ Touch Imaging System (Bio-Rad).

### Sperm immunocytochemistry

Spermatozoa were washed twice in PBS, attached onto glass coverslips and fixed with 4% paraformaldehyde (PFA) in PBS at room temperature (RT) for 10 min. The fixed samples were washed three times in PBS and permeabilized with 0.1% Triton X-100 in PBS at RT for 10 min. The permeabilized samples were blocked with 1% BSA in PBS (for CAII) or 5% BSA in PBS (for CAIV, MCT2, acetylated tubulin) at RT for 1h. Spermatozoa were stained with anti-CAII (20 µg/ml in Pierce™ Immunostain Enhancer), CAIV (10 µg/ml in Pierce™ Immunostain Enhancer), MCT2 (5 µg/ml in 5% BSA/PBS), CATSPER1 (10 µg/ml, in 5% BSA/PBS) and acetylated tubulin (2 µg/ml, in 5% BSA/PBS) at 4°C overnight. After washing in PBS, the samples were incubated with corresponding secondary antibody (1:1,000). Immunostained samples were mounted with Prolong Gold (Invitrogen) and cured for 24h before imaging.

### Confocal and 3D structured illumination microscopy (SIM) imaging

Confocal and SIM imaging was performed as previously described ^56,61^. Briefly, cured samples were used for confocal imaging performed with either a Zeiss LSM710 Elyra P1 using a Plan-Apochromat 63X/1.40 and an alpha Plan-APO 100X/1.46 oil objective lens (Carl Zeiss) or a Nikon Eclipse Ni-E equipped with a DS-Ri2 camera using a Plan-APO 40X/0.95. A 488 nm laser (200 mW) was used for Alexa 488 (Invitrogen), 561 nm laser (200 mW) was used for Alexa 568 (Invitrogen), and 642nm laser (150 mW) was used for Alexa 647 (Invitrogen) for SIM imaging. Raw image processing and rendering was performed using Zen 2012 SP2 software (Carl Zeiss).

### pH determination

#### Live animal epididymal and vas deferens luminal pH determination

After anesthesia with Nembutal (McKesson, 60mg/kg body, i.p.), mice were placed on a heating pad to measure the luminal pH of cauda epididymis and vas deferens. The vas deferens was carefully incised, taking care not to damage the deferent artery, and the fluid was expressed with forceps. For cauda epididymis, a similar approach was used, with an incision made in the distal cauda to collect fluid. A droplet of the collected fluid was applied directly to a pH strip (Hydrion, Micro Essential Lab) and pH was read visually by three independent observers. Analysis was performed on three WT, *Car2^-/-^*, and *Car4^-/-^* mice, using two cauda and two vas deferens from each mouse. pH measurements were performed in triplicate for each sample. At the end of the experiment, mice were euthanized by cervical dislocation.

#### Single live sperm pH_i_ imaging

For single-cell fluorometry, time-lapse images were acquired using an Axio observer Z1 microscope (Carl Zeiss) equipped with a pco.edge cMOS camera and DG-4. The fluorescence signal was excited at 560 nm (543/22 nm filter) and emitted at 585 nm (568LP nm filter). To allow a direct comparison of intrinsic pH-regulatory capacity, pH_i_ measurements were performed in a standardized HS buffer (pH 7.4). 1×10^6^ non-capacitated or capacitated spermatozoa from the cauda epididymis or vas deferens in 500 µl HS buffer were loaded with 5 µM pHrodo-Red-AM (Thermo Fisher) and 0.05% Pluronic F-127 in the dark at 37°C for 30 min. Spermatozoa were attached to a Delta T chamber coated with fibronectin (0.1 mg/ml) for 15 min at 37°C. The staining solution was replaced with 500 µl HS buffer (pH 7.4). The basal pH_i_ of only motile spermatozoa with their head attached to the surface was recorded for one minute. ROIs were drawn restricted to the sperm head. After one minute, the HS buffer (pH 7.4) was exchanged to HS (pH 7.4) supplemented with 15 mM NH_4_Cl to induce intracellular alkalinization, followed by a buffer change back to HS (pH 7.4). For pH calibration, HS buffers containing 15 µM nigericin adjusted to pH 5.4, 6.4, 7.4 and 8.4 with HCl or NaOH were used. Calculated standard curve was used to determine pH_i_ of wt and *KO* sperm. The percentage change in pH_i_ (ΔpH_i_ %) was calculated by setting the non-capacitated wt pH_i_ to 100 % and calculating the percentage change in pH_i_ of the non-capacitated and capacitated *KO* sperm, as well as the capacitated wt sperm. Mean ΔpH_i_ after NH_4_Cl treatment was calculated by mean pH_NH4Cl_ / mean pH_Basal_ for each spermatozoon.

#### pH_i_ determination of sperm populations using a fluorescence plate reader

A fluorescence plate reader (Infinite^®^ 200 PRO, Tecan) was used to determine pH_i_ of sperm suspension. To allow a direct comparison of intrinsic pH-regulatory capacity, pH_i_ measurements were performed in a standardized HS buffer (pH 7.4). Briefly, non-capacitated or capacitated spermatozoa from the cauda epididymis (3×10^6^ cells/ml) were loaded with 5 µM pHrodo-Red-AM (Thermo Fisher) and 0.05% Pluronic F-127 in 1 ml HS buffer in the dark at 37°C for 30 min. The staining solution was replaced by HS buffer (pH 7.4) after one centrifugation step (5 min, 300 g). 4×10^5^ cells/200 µl were added to each well and triplicates of each genotype were analyzed. After measuring basal pH_i_, NH_4_Cl (final concentration: 15 mM) was added to each well using a multi-channel pipette and the change in pH_i_ was recorded. For pH calibration, 200 µl of HS buffer containing 15 µM nigericin was adjusted to either pH 5.4, 6.4, 7.4 or 8.4 with HCl or NaOH. Errors due to dye loading or possible differences in sperm concentration were avoided by constructing an internal calibration curve for each measurement, as is done for other assays. Three wells of HS-buffer with unloaded sperm were measured as a background signal control. Reading was performed at 37°C using an excitation wavelength of 560 nm and an emission wavelength of 585 nm. The percentage change in pH_i_ (ΔpH_i_ %) was calculated by setting the non-capacitated wt pH_i_ to 100 % and calculating the percentage change in pH_i_ of the non-capacitated and capacitated *KO* sperm, as well as the capacitated wt sperm. Mean ΔpH_i_ after NH_4_Cl treatment was calculated by mean pH_NH4Cl_ / mean pH_Basal_ for each sperm population.

#### Single live sperm Ca^2+^ imaging

For Ca^2+^ Imaging, 2×10^6^ non-capacitated caudal sperm were loaded with 4 µM Fluo4-AM and 0.05% Pluronic F-127 in 500 µl HS-buffer for 30 min at 37°C in the dark and washed one time in PBS by centrifugation (3 min, 500 g). Fluo4-loaded 2×10^5^ cells in 250 µl HS buffer were plated onto a Delta T chamber treated with Poly-L-Lysine (0.1 mg/ml) for 15 min. The HS buffer was replaced with 500 µl of fresh HS buffer to remove unbound spermatozoa. Time-lapse images were captured using a 60x objective of an Axio observer Z1 microscope (Carl Zeiss) equipped with a pco.edge cMOS camera. First, the basal fluorescence intensity (t_0 s_ - t_60 s_) was recorded. The sperm cells were then depolarized by adding a 10 µl drop of K8.6 solution directly onto the sperm focused in the field of view for HS to reach final 138 mM KCl and 20 mM NH_4_Cl. Depolarization was recorded for additional 60 s (t_60 s_ - t_120 s_). Finally, 5 µl of Ionomycin (100x; final concentration= 10 µM) was pipetted onto the same sperm and the fluorescence signal was recorded for a further 60 s (t_120 s_ – t_180 s_). Ionomycin saturates the internal dye with Ca^2+^ and therefore serves as a positive control. ΔF was calculated for each sperm by the equation mean F_K8.6_ - mean F_Basal_, where F_K8.6_ represents the fluorescence intensity after K8.6 addition, F_Basal_ the resting fluorescence. Values were normalized to the mean ΔF/F_0_ of the wt.

#### 2D Sperm motility analysis

For 2D sperm motility analysis, sperm suspension was placed in a pre-warmed 20 µm deep chamber slide (Leja products B.V.) and motility was examined at 37°C using a CASA system (IVOS^®^ II, Hamilton Thorne INC.). Images were acquired at 60 Hz. CASA was used to measure total motility (%) and curvilinear velocity (VCL) (µm/s). At least 200 motile spermatozoa were analyzed for each experiment, with 3 biological replicates. Sperm concentration (sperm/ml) was determined by using a Hemocytometer counting chamber (Neubauer).

#### 4D sperm motility analysis using Digital Holographic Microscopy

The implementation of four-dimensional (4D) motility analysis using digital holographic microscopy (DHM) is described elsewhere. Briefly, the imaging was conducted using a DHM^TM^ T-1000 (Lyncée Tec SA) operating in off-axis transmission mode, equipped with a 666 nm laser diode source, a 20×/0.4 NA objective, and a Basler aca1920-155um CCD camera (Basler AG). All experiments were performed at 37°C using a 100 µm deep chamber slide (Leja products B.V.) and a frame rate of 100 Hz. Off-line processing of the stored holographic images and subsequent tracking of sperm heads to obtain X-, Y-, and Z-plane coordinates for the entire trajectory was performed using the supplied Koala (Ver. 6; Lyncée Tec SA) and open-source Spyder (Python 3.6.9) software. Sperm motility parameters including three-dimensional (3D) curvilinear velocity (VCL, in μm/s) and two-dimensional (2D) amplitude of lateral head displacement (ALH, in μm), were calculated using the obtained X, Y, and Z plane coordinates. Frame-by-frame tracing of flagellar images in stacks of reconstructed XY projections (8-bit TIFF format, 100 fps, 10-frame time) was conducted using a macro written in Igor Pro™ Ver. 6.36 (Wavemetrics) to determine X, Y, and Z values for sperm flagellar motion. ImageJ V1.50i (National Institutes of Health) was used to adjust the brightness and contrast of the reconstructed XY projections (resolution of 800 x 800 pixels). Z-plane coordinates were calculated from the received X, Y coordinates of the flagellar tracks and a script written in Spyder and Koala. A P/U value (3.7466), which is defined as the quotient of the objective magnification (x20) and the pixel size (5.34 μm) of the camera (Basler aca1920–155 μm), was used to convert pixels to microns. The Z-plane data was smoothed using a seventh-order polynomial in Igor Pro™. For 4D sperm motility analysis, 1×10^6^ sperm/ml were loaded onto the deep chamber slide and 15 free-swimming single murine and 30 human sperm were analyzed using three male mice from each genotype (wt, *Car2^-/-^*, *Car4^-/-^*) or 6 donors.

## Statistic

Statistical analysis was performed with GraphPad Prism (Vers.10, Statcon GmbH). One-way, two-way analysis of variance (ANOVA) and Student’s *t*-test were used to calculate the significance in differences of mean or median values. Differences were classified as significant at p <.05. Numerical results are presented as medians and interquartile ranges or as mean ± SEM. N = number of independent experiments.

## Supporting information

Wiesehoefer et al_Supplementary Materials

## Supplementary Materials

Figs. S1 to S5

Table S1 to S2

## Acknowledgement

We thank Sara F. Finnegan and Tianyu Zhu for their help with managing the *Car2*^-/-^, *Car4*^-/-^, *Slo3*^-/-^, and *Catsper1*^-/-^ mouse lines, Katerina Politi, from the Department of Pathology at the Yale School of Medicine, for our use of Tecan Infinite 200 Pro plate reader, Natalie Knipp and Jaroslaw Dankert for their help with managing mice at the Department of Anatomy at the University Clinic Essen.

## Author contributions

**Caroline Wiesehöfer:** Conceptualization; methodology; formal analysis; data curation; writing - original draft; writing – review and editing

**Cameron C. Gardner:** formal analysis; data curation; writing – review and editing

**Aura Stoskus:** Formal analysis; data curation

**M. Agustina Battistone:** Methodology; data curation; writing – review and editing; resource; funding acquisition

**Gunther Wennemuth:** Conceptualization; resource; supervision; funding acquisition

**Jean-Ju Chung:** Conceptualization; writing - original draft; writing – review and editing; resource; supervision; funding acquisition

## Data and materials availability

Raw data associated with Figures have been deposited at Open Science Framework. All other data needed to evaluate the conclusions in the paper are present in the paper of the Supplementary Materials. Yale University requires an MTA for new reagents generated in the study.

