## Supplementary material for "Coordinated CAII and CAIV establish a dual pH-regulatory axis essential for CatSper activation during sperm maturation and capacitation": Wiesehoefer et al_Supplementary Materials

The PDF contains Supplemental Figures 1-5 and Supplemental Tables 1-2.

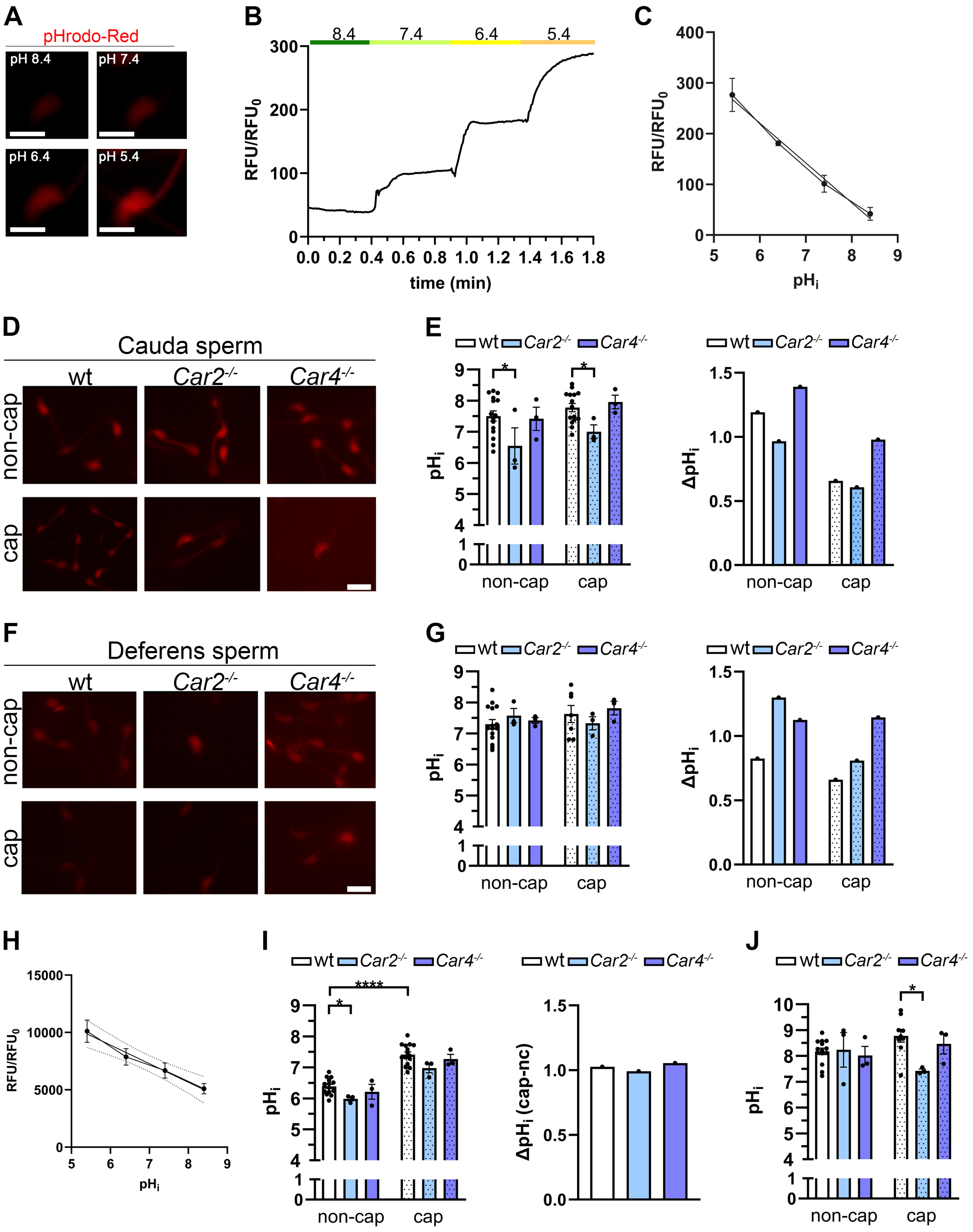


**Supplemental Fig. 1: CAII deficiency, but not CAIV deficiency, impairs internal alkaline challenge.**
(A-C) Calibration of the pH-sensitive dye pHrodo-Red using non-capacitated cauda spermatozoa. (A) Representative fluorescence images of sperm heads incubated in buffers of varying pH in the presence of 15 μM Nigericin. Scale bar= 5 μm. (B) Corresponding relative fluorescence intensity. (C) Standard calibration curve correlating relative fluorescence units (RFU) to $pH_i$. (N=3). (D) Representative

fluorescence images of non-capacitated (upper part) or capacitated (lower part) wt, *Car2*$^{-/-}$ and *Car4*$^{-/-}$ sperm heads of sperm isolated from the cauda epididymis. Scale bar= 10 µm (E) $pH_i$ quantification after challenging non-capacitated (open bars) and capacitated (dotted bars) sperm from the cauda epididymis with 15 mM $NH_4Cl$, derived from traces as shown in (Fig. 2A). Sample sizes are indicated as N = number of mice and n= number of individual cells analyzed. Non-capacitated $N_{wt}$=15, $N_{KOs}$=3, n≥21, capacitated $N_{wt}$=15, $N_{KOs}$=3, n≥14. (F) Representative fluorescence images of non-capacitated (*upper*) or capacitated (lower) wt, *Car2*$^{-/-}$ and *Car4*$^{-/-}$ sperm heads of sperm isolated from the vas deferens. Scale bar= 10 µm (G) Net $pH_i$ increase ($\Delta pH_i$) after 15 mM $NH_4Cl$ addition for non-capacitated (open bars) and capacitated (dotted bars) cauda sperm. $\Delta pH_i$ was calculated for each genotype by subtracting the mean non-capacitated or capacitated $pH_i$ from the mean non-capacitated or capacitated $NH_4Cl$ $pH_i$ shown in panel D. (H-J) Multi-cell pH analysis of motile cauda sperm. Representative standard curve based on the pH-dependent pHrodo-Red fluorescence units (RFU) to $pH_i$ (H). $pH_i$ determination of pHrodo-Red loaded non-capacitated (empty bar) and capacitated (dotted bar) wt (white), *Car2*$^{-/-}$ (blue) and *Car4*$^{-/-}$ (violet) cauda sperm (I). Standard curve used in analysis (*left*). Net $pH_i$ increase ($\Delta pH_i$) during capacitation for sperm from the cauda epididymis (*right*). $\Delta pH_i$ was calculated for each genotype by subtracting the mean non-capacitated $pH_i$ from the mean capacitated $pH_i$ shown in panel G. $pH_i$ quantification after 15 mM $NH_4Cl$ treatment (J). Data are shown as mean ± SEM. Sample sizes are indicated as N = number of mice and n= number of individual cells analyzed. $N_{wt}$=15, $N_{KOs}$=3; n= $3x10^6$ cells/ml (E, G). Non-capacitated: $N_{wt}$=14, $N_{KOs}$=3, capacitated: $N_{wt}$=12, $N_{KOs}$=3; n= $3x10^6$ cells/ml (I). Statistical significance was determined by unpaired Students t-test. *p ≤ 0.05, ****p ≤ 0.0001. *Related to* Figure 2.

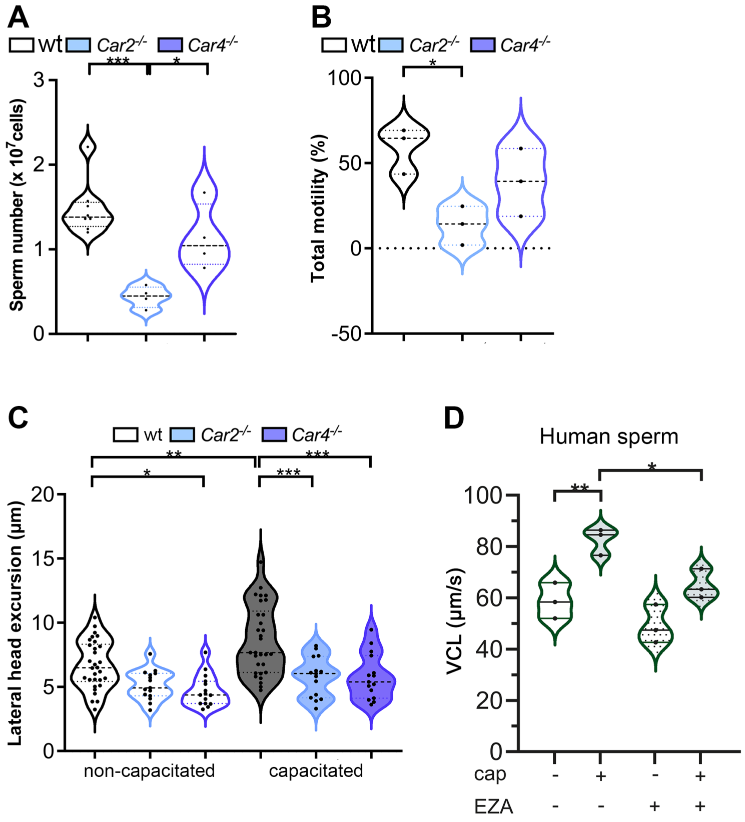


**Supplemental Fig. 2. Hyperactivated motility is compromised in sperm from *Car2* and *Car4*-null mice.** (A) Sperm number determination using a Hemocytometer. (B) 2D motility analysis reals a decreased total motility of *Car2*$^{-/-}$ (blue), and *Car4*$^{-/-}$ (violet) sperm. (C) Quantification of the lateral head excursion analyzed by DHM of non-capacitated (open violins) and capacitated (filled violins) sperm. Each point represents a single sperm (n > 15 per condition). (D) Effect of the pan-carbonic anhydrase inhibitor ethoxyzolamide (EZA, 5 µM) on human sperm motility analyzed by CASA. Data are from N=3 donors, with n>600 cells. Data are presented showing the median (thick dash lines) and interquartile range (thin dash lines). Asterisk indicates a significant difference from wt under the same condition: *$p \leq 0$ .05, **$p \leq 0.01$, ***$p \leq 0.001$. *Related to* Figure 3.

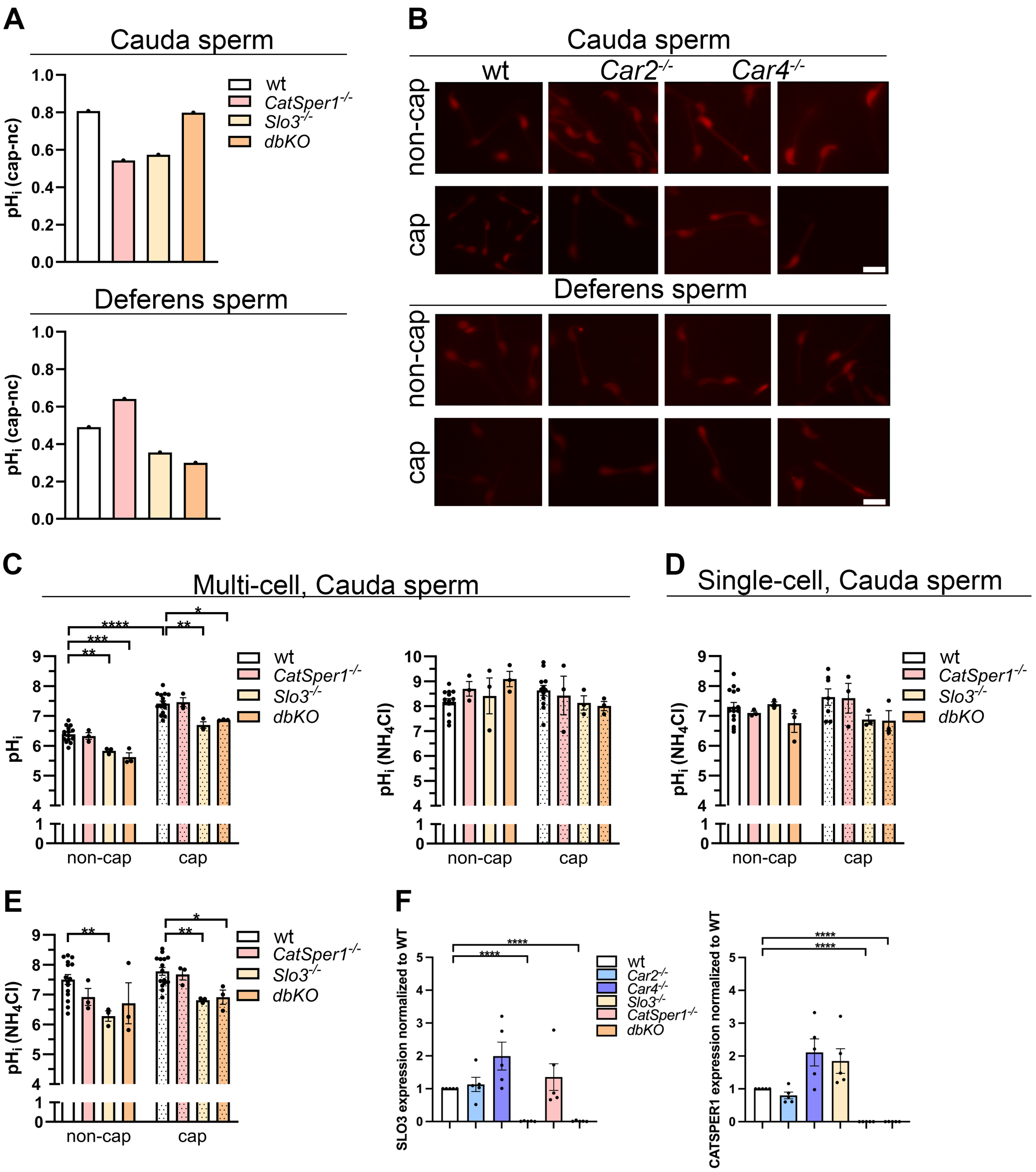


**Supplemental Fig. 3. *Slo3* deficiency causes acidification in the sperm.** (A) Single-cell pH-imaging of spermatozoa. Capacitating conditions induce alkalinization in all analyzed sperm isolated from the cauda (*upper diagram)* or vas deferens (*lower diagram*) *of* wt (black), *Catsper1*$^{-/-}$ (pink), *Slo3*$^{-/-}$ (beige), and *Slo3*$^{-/-}$*, Catsper1*$^{-/-}$ double knock-out (*dbKO*, orange) males. (B) Representative fluorescence images of pHRodo-loaded non-capacitated and capacitated murine sperm isolated from the cauda (*upper part*) or the vas deferens (*lower part*) from (A). Scale bar = 10 µm. (C) Multi-cell pH analysis of motile cauda sperm by fluorescent plate reader. Basal $pH_i$ (left) and $pH_i$ determination after 15 mM $NH_4Cl$ contact (right) of pHrodo-Red loaded non-capacitated (empty bar) and capacitated (dotted bar) sperm as described above using analysis specific calibration curve of Supplemental Fig. 1F. (D-E) Single-cell $pH_i$ analysis after $NH_4Cl$ treatment of cauda (D) and vas deferens (E) sperm calculated from fluorescence intensity and calibration curve from Supplemental Fig. 1C. (F) Densitometric quantification of SLO3 (left) and CATSPER (right) protein bands from the immunoblots (Fig. 4E). Data in (C, D, E) are presented as mean ± SEM. Sample size for multi-cell $pH_i$ experiments (C): $N_{wt}$=15, $N_{KOs}$=3; n= 3x$10^6$ cells/ml; for single-cell $pH_i$ experiments (D-E): N=3-15 mice per genotype, n≥23 cells per experiment. For protein quantification (F), each point represents a biological replicate from an individual male. Statistical significance was determined by unpaired Students t-test. *p ≤ 0.05, **p ≤ 0.01, ***p ≤ 0.001, ****p ≤ 0.0001. *Related to* Figure 4.

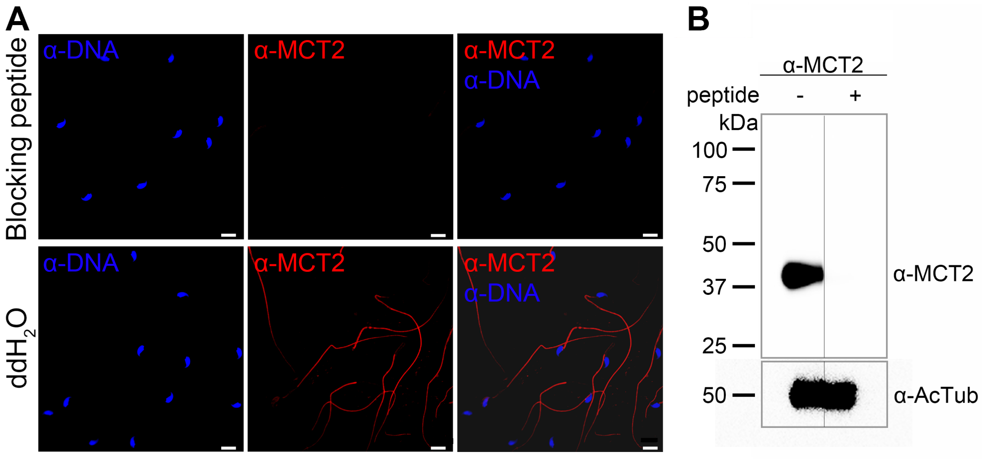


**Supplemental Fig. 4. Anti-MCT2 antibody specificity validation.** (A) Immunocytochemistry and (B) immunoblot performed with anti-MCT2 antibody either preabsorbed with the MCT2 peptide used to generate anti-MCT2 antibody (blocking peptide) or preabsorbed with vehicle (dd$H_2O$). (A) MCT2 is stained in red and nuclei stained blue. Scale bar = 10 µm. *Related to* Figure 5.

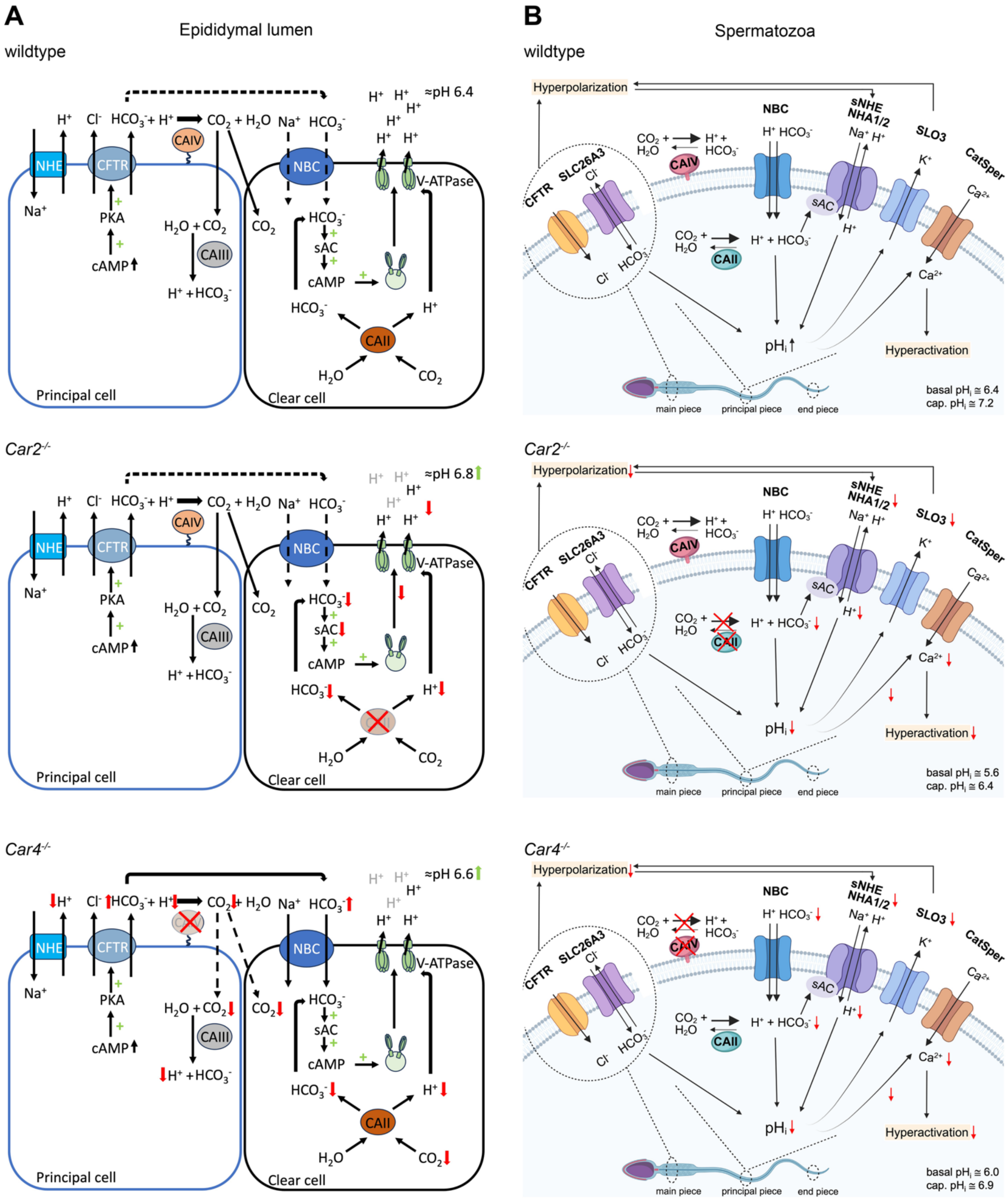


**Supplemental Fig. 5. Putative mechanism of pH and HCO3$^-$ regulation by CAII and CAIV in the epididymides and murine spermatozoa.** (A-B) pH regulation in the epididymis (A) and spermatozoa (B) of wt (first), *Car2*$^{-/-}$ (second) and *Car4*$^{-/-}$ (third) mice. CFTR: cystic fibrosis transmembrane regulator; SLC26A3: chloride-bicarbonate exchanger; sNHE: sperm-specific sodium-proton exchanger (SLC9C1 family); NHA1/2: sodium-proton antiporters (SLC9B family); NBC: sodium-bicarbonate co-transporter; SLO3: sperm-specific potassium channel; CatSper: pH sensitive sperm-specific $Ca^{2+}$ ion channel; sAC: soluble adenylate cyclase; CA: carbonic anhydrase. (B) Created with BioRender.com (https://app.biorender.com/illustrations/68b82299b885cdfc6533b502). *Related to* Figures 1 and 2.

**Supplemental Table 1. Source of antibodies used in the study**

| Antibodies | Source | Identifier |
|---|---|---|
| Sheep polyclonal anti-hu/mu Carbonic Anhydrase II | Invitrogen | Cat# PA5-33167; RRID: AB_2550596 |
| Goat polyclonal anti- mu Carbonic Anhydrase IV | Thermo Fisher Scientific | Cat# PA5-47312; RRID: AB_2607329 |
| monoclonal anti-pY (clone 4G10) | Cell Signaling Technologies | Cat# #96215 |
| Mouse monoclonal anti-SLO3 (clone S2-16) | Novus Bio | NBP1-44982 |
| Mouse monoclonal anti-Acetylated Tubulin (clone 6-11B-1) | Sigma-Aldrich | Cat# T7451 |
| Rabbit polyclonal anti-MCT2 | This study | |

**Supplemental Table 2. Sources of reagents used in the study**

| Reagents | Source | Identifier |
|---|---|---|
| Dimethyl Sulfoxide, DMSO | AmericanBio | Cat# AB03091 |
| Dithiothreitol, DTT | Bio Rad | Cat# 1610610 |
| EmbryoMax M2 Medium (1X), Liquid, with phenol red | EMD Millipore | Cat# MR-015-D |
| EmbryoMax Human Tubal Fluid (HTF) (1X), liquid, for Mouse IVF | EMD Millipore | Cat# MR-070-D |
| ethoxzolamide (EZA) | TargetMol | Cat# T5012 |
| Fibronectin, Human Plasma | Sigma-Aldrich | Cat# 341635 |
| Fluo4-AM | Thermo Fisher | Cat# F14201 |
| Ionomycin | Thermo Fisher | Cat# I24222 |
| Nigericin | Sigma-Aldrich | Cat# SML1779 |
| pHrodo-Red-AM | Thermo Fisher | Cat# P35372 |
| Pierce™ Immunostain Enhancer | Thermo Scientific | Cat# 46644 |
| Pluronic F-127 | Invitrogen | Cat# P3000MP |
| Poly-L-Lysine solution, 0.1% (w/v) | Sigma-Aldrich | Cat# P8920 |
| Triton X-100 | Sigma-Aldrich | Cat# T8787 |
| 4xLDS sample buffer | GenScript | Cat# M00676-10 |
